# Methods for high throughput discovery of fluoroprobes that recognize tau fibril polymorphs

**DOI:** 10.1101/2024.09.02.610853

**Authors:** Emma C. Carroll, Hyunjun Yang, Julia G. Jones, Abby Oehler, Annemarie F. Charvat, Kelly M. Montgomery, Anthony Yung, Zoe Millbern, Nelson R. Vinueza, William F. DeGrado, Daniel A. Mordes, Carlo Condello, Jason E. Gestwicki

## Abstract

Aggregation of microtubule-associated protein tau (MAPT/tau) into conformationally distinct fibrils underpins neurodegenerative tauopathies. Fluorescent probes (fluoroprobes), such as thioflavin T (ThT), have been essential tools for studying tau aggregation; however, most of them do not discriminate between amyloid fibril conformations (polymorphs). This gap is due, in part, to a lack of high-throughput methods for screening large, diverse chemical collections. Here, we leverage advances in protein adaptive differential scanning fluorimetry (paDSF) to screen the Aurora collection of 300+ fluorescent dyes against multiple synthetic tau fibril polymorphs. This screen, coupled with orthogonal secondary assays, revealed pan-fibril binding chemotypes, as well as fluoroprobes selective for subsets of fibrils. One fluoroprobe recognized tau pathology in *ex vivo* brain slices from Alzheimer’s disease patients. We propose that these scaffolds represent entry points for development of selective fibril ligands and, more broadly, that high throughput, fluorescence-based dye screening is a platform for their discovery.

## Introduction

Tau is an intrinsically disordered, microtubule-binding protein that assembles into β-sheet rich fibrils within the neurons of patients suffering from a family of devastating neurodegenerative diseases, known as tauopathies^1–3^. In Alzheimer’s disease (AD), one of the most common tauopathies, these tau fibrils are characterized as either straight filaments (SFs) or paired helical filaments (PHFs)^4^, suggesting that the same protein might be able to form fibrils with distinct molecular structures. Indeed, cryo-electron microscopy experiments have revealed that the core structure of patient-derived tau fibrils adopts different molecular conformations or folds^5^, referred to here as polymorphs. Interestingly, these fibril polymorphs appear to be disease-specific; tau fibril structures differ between some clinically distinct tauopathies but are recapitulated in patients with the same disease^6^. Together, these observations have driven interest in the development of optical reagents that can rapidly discriminate between tau fibril polymorphs.

Organic dyes have been essential tools for studying amyloid fibrils for many decades^7, 8^. The power of these reagents is that their spectral properties change when they are bound to fibrils, making them relatively straightforward to use. For example, the most widely used fluorescent probe (fluoroprobe) is thioflavin T (ThT) and its fluorescence intensity is dramatically increased when bound to amyloids^7, 9–12^. Likewise, fluoroprobes based on Congo Red, curcumin, polythiophenes and other scaffolds^13–18^ have proven to be convenient tools for studying how fibrils form *in vitro* and in cells and tissues. Fluoroprobes have also been used as competitors to identify non-fluorescent compounds by displacement^19^. While fluoroprobes have played critical roles in studying tauopathies, they typically lack specificity for different fibril polymorphs^20, 21^. Indeed, their generality is often a great strength because a single fluoroprobe, such as ThT, has the versatility to detect a wide range of fibrils, largely independent of sequence or substructures. Yet, the field would also benefit from complementary fluoroprobes that are selective for subsets of tau polymorphs.

Most amyloid ligands have been generated by creating close structural analogs of established amyloid-binding scaffolds, such as ThT or curcumin^14, 22, 23^. While those efforts are often successful in producing analogs with improved properties, such as brightness or permeability, they do not typically involve sampling of a wide range of chemical space. We hypothesized that more diverse starting points, with a better chance of being selective, might be uncovered by screening dye collections that contain a variety of chemical scaffolds. We saw an opportunity to address this persistent challenge in the recent development of a protein adaptative differential scanning fluorimetry (paDSF)-based platform that leverages the Aurora collection of 300+ chemically diverse dyes (**Extended Data 1**)^24^. To test this idea, we first needed to produce large quantities of tau fibrils (*e.g.,* micrograms to milligrams) with different structures. Towards that goal, we turned to the P301S point mutation in *MAPT*. Briefly, this mutation is linked to frontotemporal dementia (FTD) and our data, and the work of others^25–28^, has shown that tau containing the P301S (or the related P301L) mutation leads to fibril structures that are distinct from those created with wild type (WT) tau protein. Thus, we reasoned that we could use synthetic fibrils composed of WT or P301S tau to sample distinct polymorphs. To further diversify the structure(s) of these fibrils, we also varied the polyanion used to induce *in vitro* tau aggregation reactions, as it has recently been shown that the identity of the polyanion inducer also contributes to fibril structure^29–31^. Then, we screened each of these fibril samples against the Aurora collection using fluorescence-based paDSF in 384-well plates and validated the resulting dye “hits” using two orthogonal secondary assays: multidimensional spectral confocal microscopy and kinetic aggregation assays. Using this workflow, we found that a subset of the “hit” molecules bound most of the tau polymorphs (*e.g.,* pan-fibril binders), while others were relatively specific to subsets of fibril conformers (*e.g.,* selective fibril binders). These molecules included compounds with coumarin and polymethine scaffolds, chemotypes that are under-represented in the field of amyloid-binding dyes^32, 33^, as well as chemotypes not previously associated with amyloid recognition. To show the utility of this approach, we validated one validated “hit” molecule for histological staining of tau deposits in postmortem brain samples from AD patients. Finally, a proof-of-concept medicinal chemistry campaign revealed initial structure-activity relationships (SAR) for the coumarins, suggesting a potential path forward to their optimization. Thus, we envision that this paDSF-enabled workflow could be further adapted for discovery of fluoroprobes that recognize disease-specific fibril conformers for tau and potentially other amyloid-prone proteins.

## Results

### Creation of a test collection of tau fibril polymorphs for initial screening

Our goal was to establish a workflow for rapid discovery of conformationally selective, fibril-binding fluoroprobes (**Fig 1**). First, we needed to produce tau polymorphs at quantities sufficient for assay development and screening (*i.e.,* micrograms to milligrams). While fibrils isolated directly from patients with tauopathies are known to have distinct polymorphs^5^, the expected sample demands led us to consider alternative sources. Synthetic fibrils made *in vitro* from recombinant, human tau containing the disease-associated mutation, P301S, are known to have a different structure from those made with WT tau^27^. Moreover, it has also been recently shown that recombinant tau can be coaxed into distinct polymorphs by replacing the salt^34^ or polyanion^29^ component of the buffer during the aggregation reaction. For example, polyanions, such as heparin or polyphosphate, are typically used to accelerate tau fibril formation *in vitro* and the identity of the polyanion has a substantial impact on the conformation of the resulting tau fibrils, as judged by transmission electron microscopy (TEM) and limited proteolysis^29–31^. Thus, to develop a workflow for the discovery of polymorph-specific fluoroprobes, we decided to create a pilot library of synthetic tau fibrils by changing both the sequence of the recombinant tau (WT *vs.* P301S) and the polyanion used in the aggregation reactions (using 13 polyanions, favoring naturally occurring biomolecules; **Table S1**). Although we do not assume that these synthetic fibrils will necessarily have the same structure as patient-derived samples^35^, they have the practical advantages of being scalable and of potentially sampling a wide conformational space.

**Figure 1.**
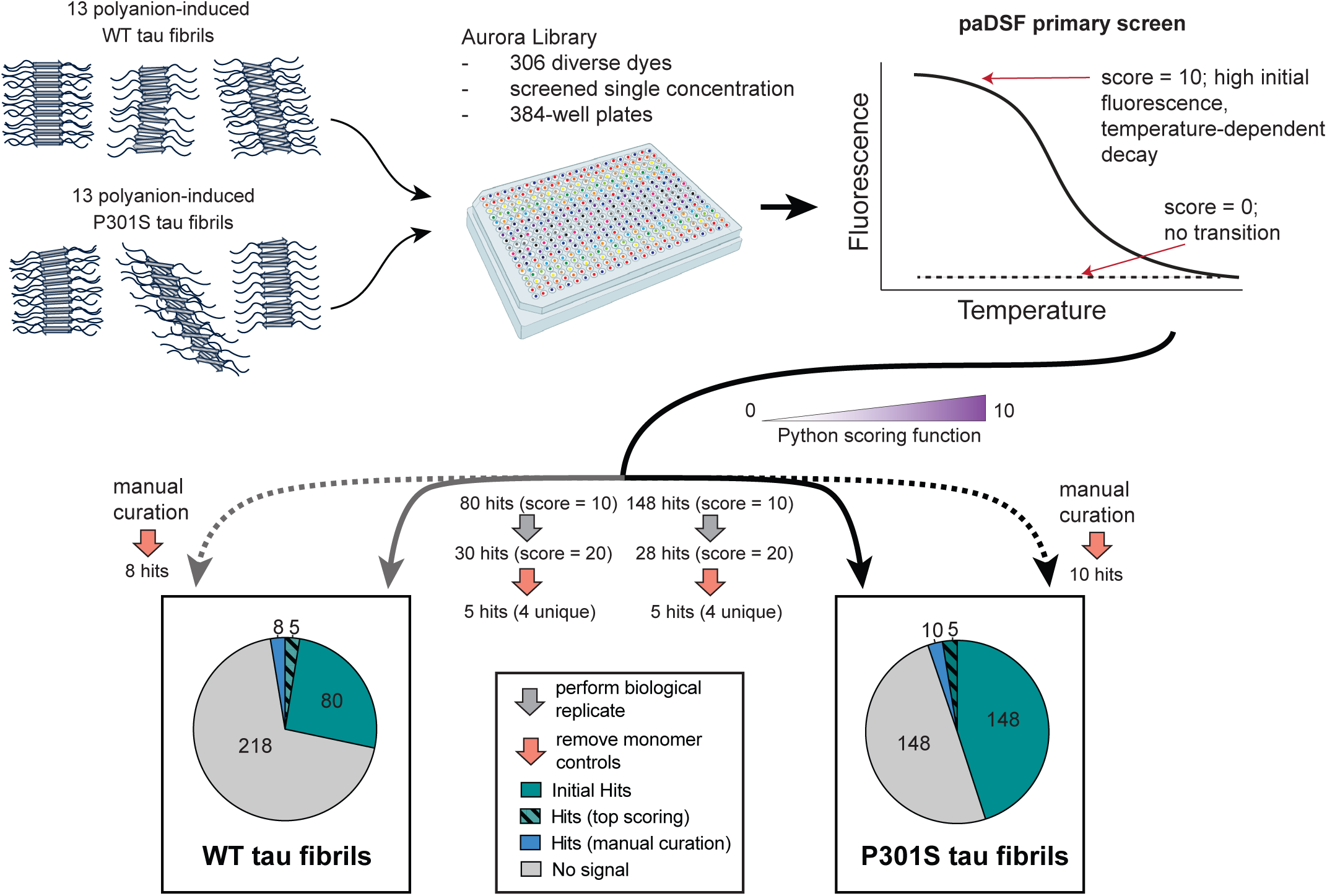
A high throughput screening platform reveals fluoroprobes that recognize tau fibril polymorphs. Schematic of the primary screening workflow and summary of the results. Synthetic tau fibrils with diverse conformations were generated using either recombinant WT or P301S tau (0N4R splice isoform), mixed with 13 different inducers (see Fig. S1 and Table S1). These 26 fibril samples were purified and incubated with the Aurora dye library in 384-well plates and then heated to generate temperature *vs.* fluorescence plots. The resulting data was scored using a Python-based function (see Methods), with the top “hits” (score = 10) being dyes with high initial fluorescence, low background in the control (polyanion inducer; no tau) and a temperature-dependent decay. The highest-scoring hits across two biological replicates (cumulative score = 20) were then compared to the second control (monomeric tau alone; salmon), yielding five hits that reacted with at least one of the WT or P301S fibrils (hashed). This list was supplemented by manual curation of other top performing dyes (blue).

With this plan in place, we expressed and purified recombinant human tau with the 0N4R splice variant, as either wild type (WT) or the P301S point mutation. These two proteins were then mixed with 13 different polyanions (**Table S1**) in aggregation reactions, yielding a total of 26 fibril samples. Each reaction was carried out for 48 hours, which was shown to be sufficient to complete fibril formation^29^, and the resulting fibrils were separated from excess reaction components (*e.g.,* unreacted monomer, soluble oligomers) by ultracentrifugation (see Methods). To validate whether some of these fibrils might have distinct structures, we resuspended a sample in buffer and treated with trypsin. The fragments remaining after proteolysis were then separated by SDS-PAGE and blotted with antibodies (named tau 13, tau 1, tau 5 and 4R) that recognize four different antigenic sites on the tau protein (**Fig S1**). This proteolysis-enabled profiling approach is commonly used to reveal which epitopes in tau are relatively included/excluded from the fibril core^27, 36, 37^ and WT 0N4R tau monomer and fibrils have already been extensively characterized by this method^29^. As a control, we confirmed that tau P301S monomer appeared at the expected molecular weight of ∼55 kDa in the absence of trypsin and that all four epitopes were completely hydrolyzed by enzyme addition (**Extended Data 2**; **Fig S1a**). As expected, fibrils formed from WT and P301S tau, both incubated with the same inducer (*i.e.,* heparin), yielded distinct patterns^27, 29^, consistent with them being different polymorphs. Similarly, varying the identity of the polyanion in the presence of P301S tau also tended to produce a variety of proteolytic patterns, which we grouped into crude categories based on qualitative measures of epitope availability (**Fig S1b**). Here, we largely focus on the differences in how WT and P301S tau fibrils bind to fluoroprobes, with the polyanion inducers serving as a secondary axis to further diversify polymorphs. Importantly, each of these fibril samples could be generated on a sufficient scale to permit downstream assay development and screening efforts.

### Development of a high throughput screening platform for detecting tau fibril fluoroprobes

In the relatively hydrophobic and rigid environment of an amyloid fibril, the fluorescence intensity of known fluoroprobes, such as ThT, is dramatically increased^7, 9–11^. Other dyes undergo a change in their fluorescence emission or excitation wavelength (*e.g.,* blue shift) in the amyloid-bound state^38^. Thus, we envisioned that fluorescence intensity and wavelength would be good surrogates for fibril binding in the protein adaptive differential scanning fluorimetry (paDSF) platform. Briefly, psDSF combines next-generation data analysis pipeline with the Aurora collection of organic dyes to probe temperature-dependent changes in the interaction(s) between dyes and proteins during heating^39^. Importantly for our purpose, the Aurora library is composed of 306 chemically diverse compounds, including those typically used in laser manufacturing, the textile industry, biological imaging and other diverse applications (for a full list, see **Extended Data 1**)^39^. This collection also features compounds from the Max A. Weaver dye collection (MWC), which has been shown to have high chemical diversity^40^. Another key feature of the paDSF platform is that it is relatively rapid: a full screen can be performed on ∼70 µg of purified protein in a standard qPCR instrument fibril and the experiment can be completed in ∼two hours. Finally, in these experiments, fluorescence is monitored in six distinct channels over a continuous temperature ramp from 25 to 95 °C, which allows the detection of changes in either wavelength and/or intensity. Thus, we envisioned combining paDSF with the 26 tau fibril samples to identify Aurora dyes that recognize fibrils (see **Fig 1**), with a special focus on those compounds that might discriminate between subsets of polymorphs (*e.g.,* WT vs P301S).

Accordingly, we aliquoted each of the 306 dyes (0.5–50 µM) into the 26 fibril fibrils samples (0.1 to 0.2 µg/mL) in 384-well plates (10 µL total volume) and performed paDSF experiments (**Fig 2**). In these screens, we expected “hits” to be fibril-dye pairs that exhibit high initial fluorescence, indicative of dye binding to the pre-formed fibril state (see **Fig 1**). We also reasoned that promising hits might display a temperature-dependent decay in fluorescence as the putative binding site(s) are impacted by heating. As controls, we counter-screened the same dyes against polyanions alone (*i.e.,* no protein) or the tau monomers alone (*i.e.,* no inducer) (**Fig S2a-d**). Together, these screens and counter-screens produced large amounts of fluorescence *vs*. time data, so we developed a Python-based scoring function to automate the analysis. Briefly, this scoring function assigns a top score of ‘10’ to those hits with the best signal-to-noise compared to the controls (see Methods and **Extended Data 3** for Python code).

**Figure 2.**
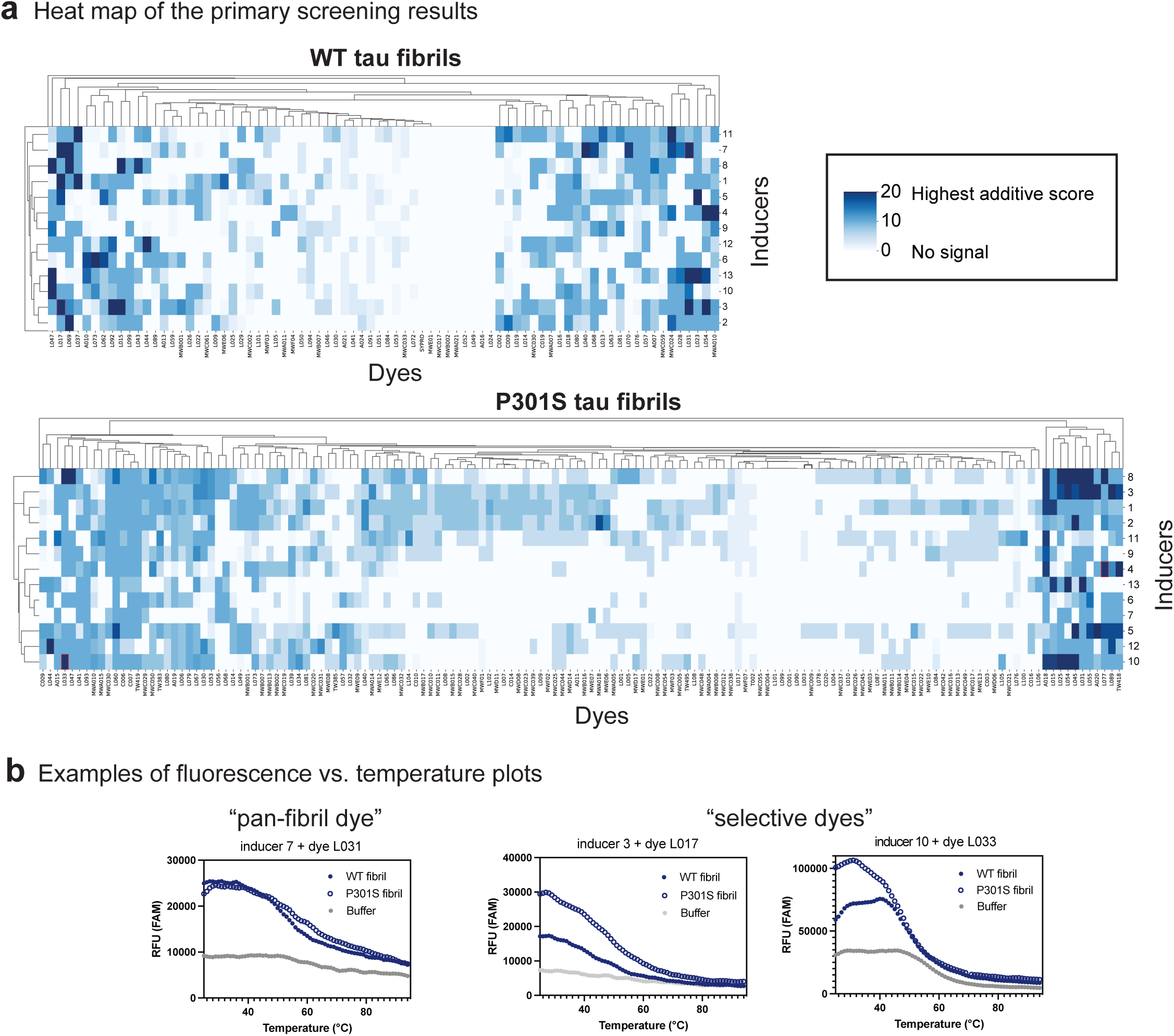
High throughput screen results uncovers fluoroprobes that interact with either WT or P301S tau fibril polymorphs. (a) Heat map of the additive scores for screens performed using WT (top) or P301S (bottom) tau fibrils. Only the highest-scoring fluoroprobes (score = 20) were taken forward for validation. (b) Representative plots of relative fluorescence units (RFU, channel denoted per plot) versus temperature, highlighting the appearance of potentially pan-fibril dyes (L031) and potentially P301S-selective dyes (L017 and L033). Graphs are representative of the two biological replicates. See **Extended Data S1** for the full dataset and **Table S1** for a list of inducers.

To identify hits, we first compared the results of the fibril screens to those of the corresponding “polyanion only” controls (see **Fig 1**). Using this approach, the WT fibrils yielded 80 initial hits with a score = 10 (26%) across all the fibril samples, while screening of the P301S tau fibrils yielded 148 hits (48%). For each initial hit, we then performed biological replicates (*i.e.,* fibrils formed on different days). We considered this step to be potentially important because amyloid fibrillization reactions can sometimes yield variable or heterogeneous outcomes. Indeed, the biological replicate screens focused us on the 30 hits (9.8%) that scored a 10 in each of the two WT screens and 28 hits (9.1%) that scored a 10 in each of the P301S fibril screens (**Fig. 1** and **Fig 2a**). From these replicate hits, we then removed any dyes that also interacted with the tau monomer controls, resulting in 5 hits (1.6%) from each of the WT and P301S screens. One of the resulting dyes was shared between the screens, suggesting the potential for pan-fibril binding (see below). Interestingly, the monomer-only control allowed us to remove a number of dyes that appeared to interact with both monomeric and fibrillar tau, despite the disorder of the monomer; yet, no dyes were identified that interacted exclusively to the monomer. Then, to supplement the list of fibril-binding “hits”, we manually re-analyzed the primary screening results to identify dyes that scored just below the cutoff but had other favorable properties (*e.g.,* low intrinsic fluorescence, diverse chemical structures; see Methods). Indeed, this process yielded an additional 8 validated hits (2.6%) from the WT screening and 10 (3.3%) from the P301S screening. Put together, this primary screening, replication and triage effort yielded 27 total fluoroprobes that interacted with at least one of the tau fibrils (**Fig 1** and **Table S2**) and it seemed to include both pan-fibril binding and potentially selective molecules.

A closer inspection of the raw data supported the accuracy of the computational analysis pipeline. For example, dye L077 only bound to P301S tau fibrils and not WT fibrils, especially in the presence of polyanion 4 (**Fig. 2b**), making it an example of a fluoroprobe with promising selectivity. By contrast, some of the fluoroprobes seemed to be more general fibril binders; for example, the dye L033 bound both WT and P301S tau fibrils that had been incubated with polyanion 10, while the dye L031 bound both WT and P301S tau fibrils incubated with polyanion 7 (**Fig. 2b**). Thus, this primary screen revealed both putatively pan-specific fluoroprobes and ones that seemed relatively selective (*i.e.,* did not bind all of the fibril samples)

### Validation of selective fluoroprobes by multidimensional fluorescence imaging

To validate the hits from the paDSF screen using an orthogonal measurement, we employed our recently developed method known as Excitation Multiplexed Bright Emission Recording (EMBER) imaging, a multidimensional fluorescence spectral confocal microscopy technique that enables detection of different tau fibril strains based upon shifts in excitation and emission spectra upon dye binding^41^. First, the 27 hit fluoroprobes were tested for direct binding to a subset of fibrils via fluorescence confocal microscopy. In these pilot experiments, we used only a subset of synthetic fibrils (WT and P301S tau, incubated with 6 different inducers; see Methods) because this subset was predicted to sample all of the polymorphs that interact with the dyes. From these preliminary microscopy experiments, 10 out of the 27 dyes (37%) were found to bind at least one of the fibrils (**Fig. 3a**). Thus, the paDSF screen had an orthogonal validation rate of ∼40%, which is in the range expected for high throughput assays. Next, we subjected these 10 dyes to the more complete EMBER imaging pipeline. Briefly, the ten fluoroprobes were each mixed with the full set of 26 tau fibril samples and imaged with confocal microscopy (**Fig 3a**). For each dye-fibril pair, we collected fluorescence information for tau fibril particles (**Fig 3b**), using a white light laser to collect 128 spectral profiles across the visible spectrum (**Fig 3c**). We then segmented the tau fibril particles from the confocal images and concatenated the excitation and emission profiles in an array for principal component analysis (PCA). Using this approach, each marker on the resulting PCA plot represents a single segmented particle in a confocal image (**Fig 3d**). Next, we asked whether any of the hit dyes could discriminate between WT and P301S tau fibrils by examining the separation of the corresponding particles in the PCA plot. Specifically, to quantify the separation of the clusters between WT and P301S tau fibrils, we employed quadratic discriminant analysis on each PCA plot to calculate discrimination scores. Satisfyingly, this process identified fluoroprobes with clear ability to discriminate between the polymorphs. For example, comparing WT vs P301S tau fibril with inducer 12, we found that dye L031 yielded a clear decision boundary from the quadratic discriminant analysis (**Fig. 3e**). Next, we averaged the discrimination scores comparing WT and P301S tau fibrils across the 13 inducers to provide an overall discrimination score for each fluoroprobe. The heatmap (**Fig. 3f**) shows that the discrimination scores ranged from 93% to 66%, with L079 and L016 having the highest overall values. Based on these data, we speculate that WT and P301S tau fibrils could remain partially distinct, even if the identity of the polyanion is changed, but this hypothesis requires additional exploration. Together, these studies show that EMBER is a valuable secondary assay, which we used to confirm fluoroprobes that bind to tau fibrils and discriminate between WT and P301S polymorphs to varying extents.

**Figure 3.**
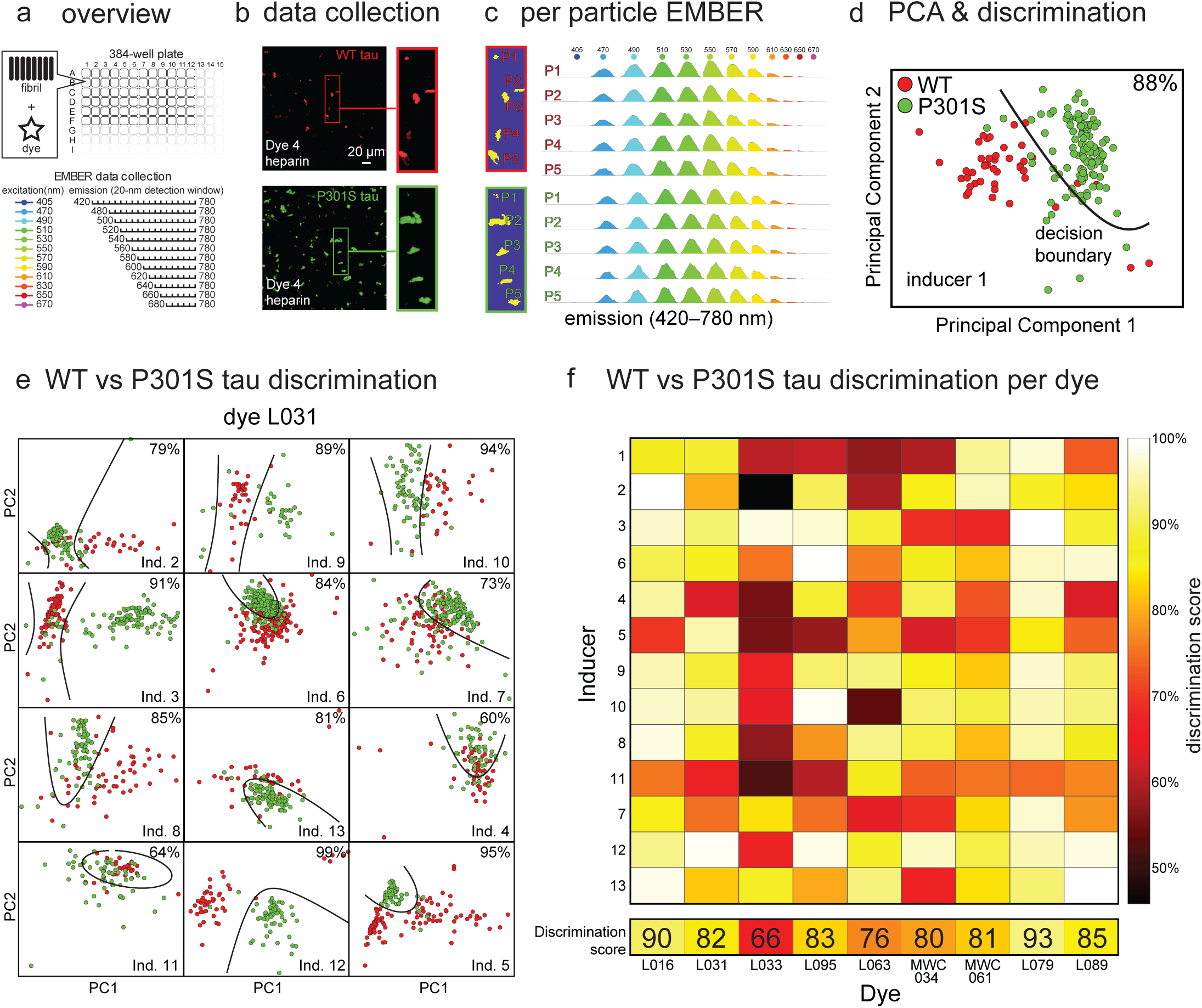
EMBER analysis suggests that fluoroprobes bind in distinct chemical environments between WT and P301S tau fibrils. (a) Overview of the EMBER (excitation multiplexed bright emission recordings) workflow. In the initial screen, the 27 hit dyes from the primary screen (see Fig 2) were screened against 12 synthetic tau fibril samples in 384-well plates. In EMBER, fluorescence data is collected at a range of excitation and emission wavelengths to explore shifts in either wavelength or intensity after dye binding to fibrils. (b) Representative EMBER results, showing individual tau fibril particles composed of either WT tau (red) or P301S tau (green). Insets show the same particles at higher resolution. This example shows WT and P301S tau fibrils with inducer 1 and dye L095. (c) For each particle, Bradley–Roth segmentation is performed across the full wavelength range to provide the EMBER plot (**Extended Data 6**). The particles are the same as in panel b. (d) Individual EMBER plots are then concatenated for principal component analysis (PCA), followed by quadratic discrimination to quantify polymorph selectivity. In this case, dye L095 was able to discriminate between WT and P301S tau fibrils (in the presence of inducer 1) with 88% accuracy. Boundaries pertaining to the fit discriminants are presented in black lines. (e) Representative data showing that dye L031 is also able to discriminate between WT and P301S tau fibril samples created using other inducers (Ind). PC1 = principal component 1. PC2 = principal component 2. (f) Discrimination heatmap for the full dataset, showing that a subset of the dyes can discriminate between tau fibril samples.

### Kinetic aggregation assays confirm the interaction(s) of fluoroprobes with tau fibrils

Together, the paDSF and EMBER screens yielded 10 validated hits that bound to tau fibrils. This list was enriched for two major chemotypes: coumarins and polymethines (**Fig. 4a**), but it was also relatively chemically diverse as evidenced by low Tanimoto pairwise similarity coefficients when compared to each other (**Fig. 4b**, **left)**. These molecules also had relatively poor Tanimoto pairwise similarity coefficients when compared to ThT (**Fig. 4b**; **right**), suggesting that they diverge from the standard fluoroprobes. Indeed, both coumarins and polymethines are relatively under-explored as amyloid ligands dyes^32, 33^ and the other compounds, MWC034 and MWC061, have never (to our knowledge) been reported to have amyloid-binding properties.

**Figure 4.**
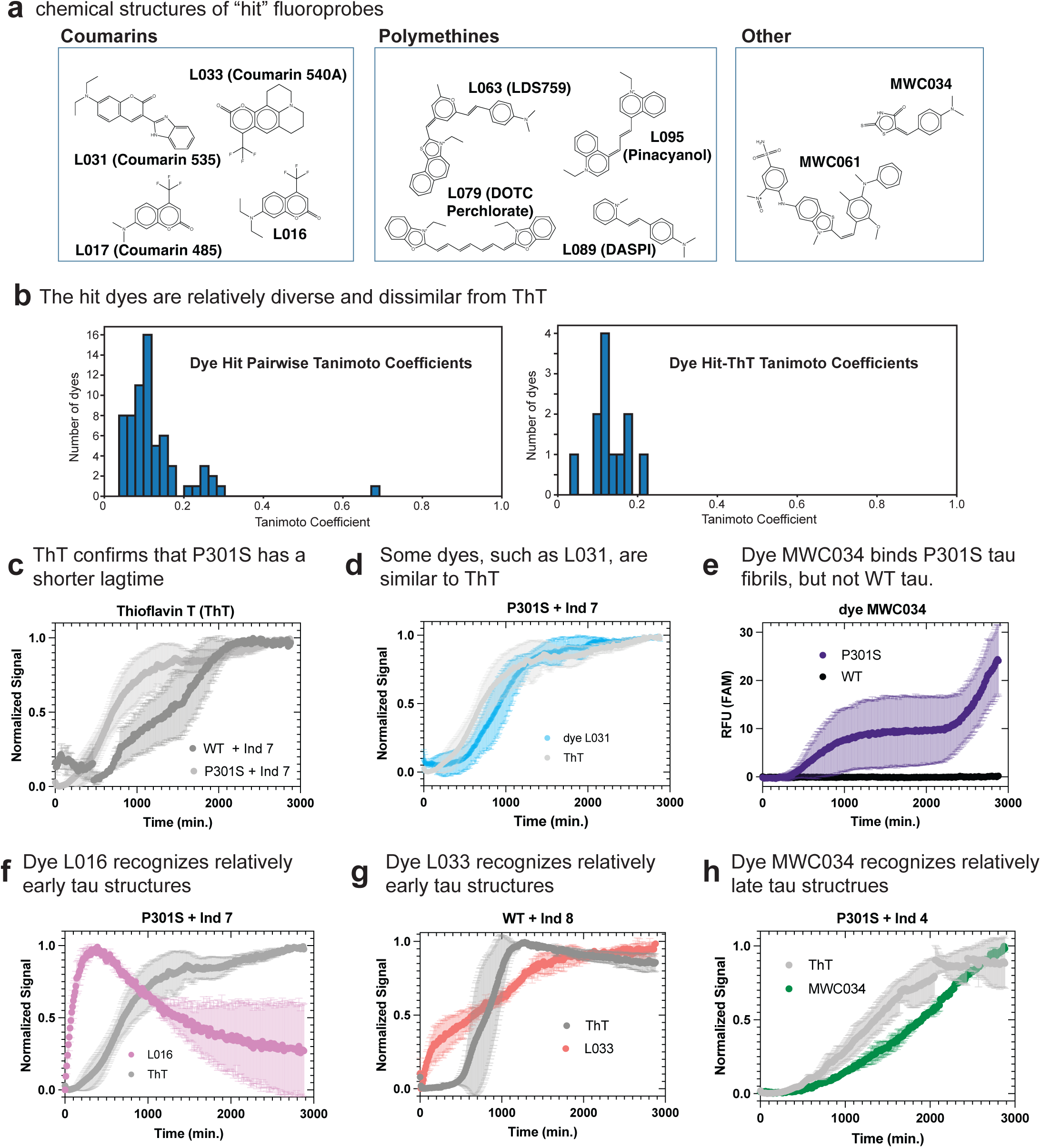
Validated fluoroprobe hits are chemically diverse and can detect tau fibril formation in real-time. (a) Chemical structures of the validated fluoroprobe hits, showing the two clusters (coumarins and polymethines). (b) Histogram of pairwise Tanimoto similarities for all hits compared with one another (left) and compared with Thioflavin T (ThT; right). Tanimoto coefficients calculated using a script created with the RDKit Python package (see **Extended Data 3**) (c-h) Kinetic aggregation assays. Either ThT or the hit dyes were mixed with WT or P301S tau and aggregation initiated with a polyanion inducer. Raw signal was normalized as a fraction of total signal to fall between 0 and 1 to facilitate comparisons. Data points are the average of replicates (n=2 or n=3) and error bars represent the standard deviation (SD). Note that only one dye was added to each sample, but the results are shown side-by-side. (c) Confirmation that P301S aggregates faster than WT, as shown using ThT and inducer 7. (d) Example of a dye, L031, that has a similar profile to ThT. Shown is P301S tau with inducer 7. (e) Example of a dye, MWC034, that only recognizes P301S tau, and not WT. Reactions contained inducer 7. (f) Example of a dye, L061, that recognizes structures early in the process than ThT. Results from P301S and inducer 7. (g) L033 recognizes relatively early structures, using WT tau and inducer 8. (h) Dye MWC034 recognizes relatively late structures, using P301S tau and inducer 4.

As a next step in validating these probes, we used them to monitor fibril formation in ‘ThT-like” kinetic aggregation assays. One goal of these experiments was to test whether any of the dyes could also be used to detect fibril formation over time. In contrast, the psDSF and EMBER platforms focus on end-stage fibrils, and we hypothesized that the binding site(s) of the fluoroprobes may appear relatively early or late in the tau aggregation reactions. In these experiments, WT or P301S tau protein was treated with eight different polyanions and the reactions shaken in an incubator in the presence of either ThT or one of the ten fluoroprobes. As expected^42^, the ThT controls produced curves for each of the aggregation reactions, with a characteristic lag time, followed by maturation of the signal until a plateau is reached (the full dataset is available in **Extended Data 4**). For example, in the presence of inducer 7, both WT and P301S tau formed ThT-active fibrils (**Fig. 4c**). These controls also confirmed that P301S tau is more aggregation-prone than WT^30, 42, 43^; the lag time was shorter for P301S tau compared to WT tau. When we replaced ThT with the “hit” fluoroprobes, nearly all of them (7/10, 70%) were active in this format, supporting the idea that the kinetic aggregation assay is another useful secondary screening platform. Moreover, we observed a range of interesting behaviors of the new dyes, allowing us to learn more about their properties. For example, the fluorescence of some compounds, such as L031, largely tracked with ThT over time (**Fig. 4d**). This result suggests that L031, like ThT, might bind a wide range of structures (*e.g.,* oligomers, fibrils), that appear during the aggregation process, at least under these conditions. Perhaps this result is not surprising because L031, like ThT, has a putative molecular rotor pharmacophore and a shared tertiary amine (**Fig. S3).** Satisfyingly, other dyes only produced a signal in the presence of P301S tau and not WT tau. For example, in the presence of inducer 7, MWC034 clearly produced a signal for P301S tau fibrils and not WT tau fibrils (**Fig. 4e**). Thus, this dye could discriminate between the two polymorphs under these conditions. Interestingly, this same dye produced a rather modest discrimination score by EMBER (76%) (see Fig 3), so it seems that the two secondary assays could emphasize distinct fluoroprobe properties. Finally, inspection of other kinetic traces suggested that some dyes, such as L016 and L033, might recognize relatively early structures in the aggregation reaction (**Fig. 4f** and **Fig. 4g**); however, this mechanistic speculation requires additional study. Likewise, the dye MWC034 seemed to detect structures that formed relatively later than ThT, at least for the P301S tau in the presence of inducer 4 (**Fig 4h**). At present, we concluded that “ThT-like” secondary assays are a relatively rapid way to validate hits from the paDSF screen and, additionally, that this step might also help categorize whether the fluoroprobe prefers binding structures that are formed relatively early or late in the fibril formation process. Taken together, this high throughput workflow, coupled with a computational analysis pipeline and two distinct secondary assays, seemed to yield multiple, promising fluoroprobe “hits”, including chemotypes that were either new or under-explored in the amyloid-binding literature.

### Fluoroprobe L095 detects tau pathology in Alzheimer’s disease patient samples

One potential use of new fluoroprobes is in the detection of tau polymorphs in brain tissue, as commonly performed with Thioflavin S^44^. Thus, we next tested whether any of our validated fluoroprobes might be useful in histopathological studies. Because we expected that some dyes may exhibit undesirable, non-specific tissue binding properties, we first performed an initial pre-screening of the 10 hit dyes in normal mouse brain slices (lacking amyloid deposits) to remove those compounds with high background fluorescence. From this screen, we selected L095 as a potentially promising candidate with low background. L095 is also known as pinacyanol and it is commonly used as a laser tuning dye and was recently evaluated as a potential amyloid ligand^45^. As a proof-of-concept, we investigated whether L095 could label tau deposits in postmortem brain samples from patients with advanced AD neuropathology (**Table S3**). In AD samples, we found that L095 successfully labeled tau neurofibrillary tangles (NFTs) with good overlap between its fluorescence signature and the staining from AT8 or pS396 antibodies that recognize pathological tau deposits (**Fig. 5a**, white arrows). Additionally, L095 labeled structures that resemble tau neuropil threads (NPTs), which were not robustly labeled by the AT8 antibody (**Fig 5a**, yellow arrow). NPTs are morphological distinct lesions of aggregated tau found within the dendritic and axonal compartments and can also be observed within dystrophic neurites, which are swollen, abnormal neuronal processes intermingled with senile plaque AD pathology^46^. Thus, this is an exciting finding, because it demonstrates the potential utility of L095 in staining a relatively understudied tau deposit.

**Figure 5.**
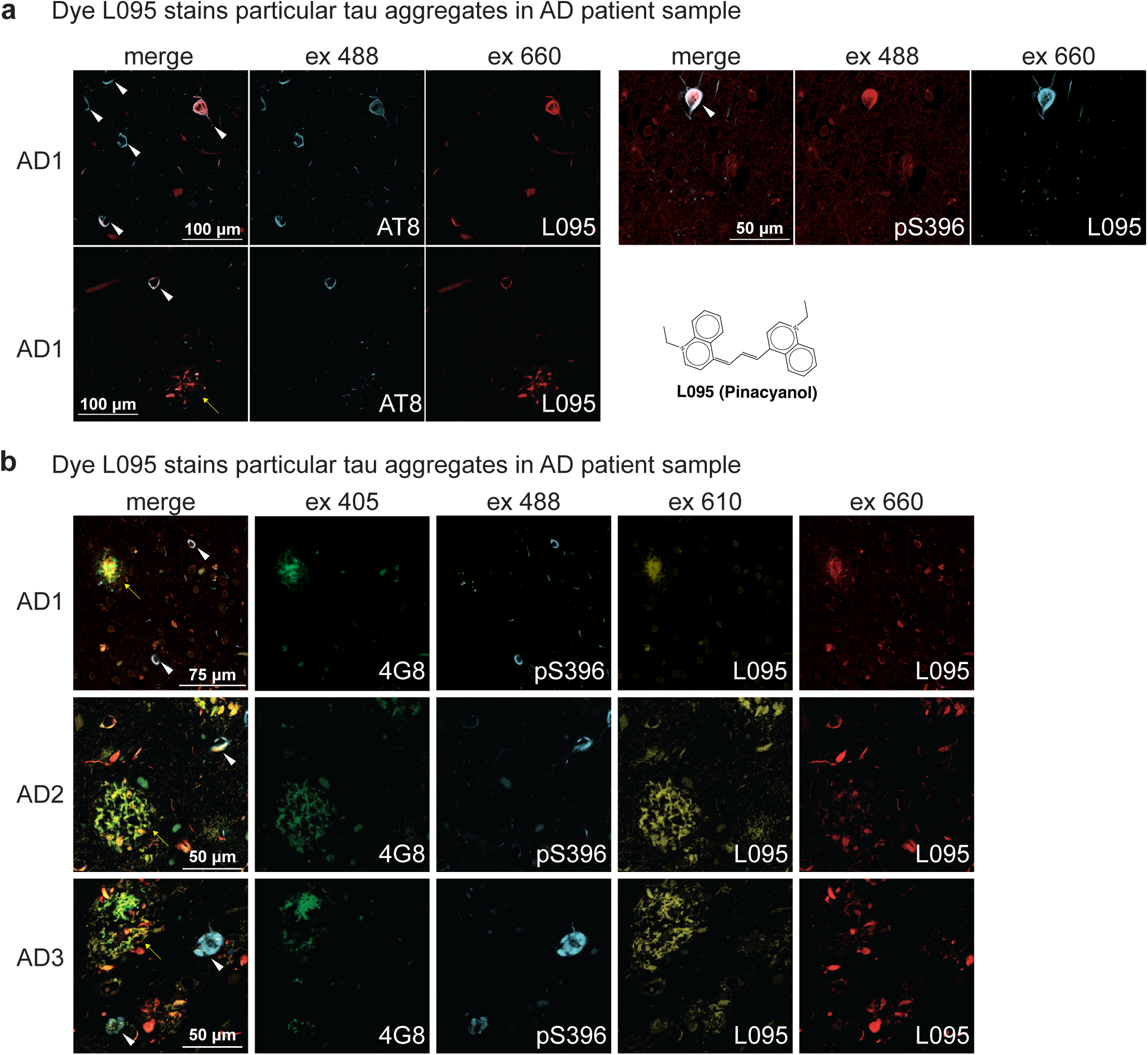
Fluoroprobe L095 recognizes tau pathology in brain tissue from human AD patients. (a) Two samples from an AD patient, labelled with the AT8 or pS396 antibody and with L095. Note that L095 only labels a subset of tau tangles (white arrows), but also labels tau pathology that is consistent with neuropil threads (yellow arrow). (b) Samples from three AD patients, stained with antibodies for either amyloid beta (4G8) or tau pathology (pS396). Note that L095 co-localizes with G48 at 610 nm (yellow arrows), but with AT8 at 660 nm (white arrows), allowing partial spectral discrimination between the two pathologies.

Since AD patient brain samples also contain deposits formed from Aβ (amyloid-β), we performed a new multi-plex staining experiment using the anti-Aβ antibody 4G8, the anti-tau pS396 antibody, and fluoroprobe L095 to understand whether the fluoroprobe might also recognize these amyloid pathologies. Indeed, we found that L095 fluorescence did overlap with both antibodies in AD samples, suggesting that it binds to both Aβ plaques and tau tangles (**Fig. 5b**). However, to be certain that L095 binds to Aβ, we confirmed its interactions in brain slices from transgenic AD mice models (APP23 and 5xFAD), which lack tau pathology, as well as its binding to synthetic Aβ40 and Aβ42 fibrils prepared *in vitro* (**Fig. S4**). While this binding to Aβ deposits was unexpected, we noted during these experiments that the fluorescence signal of L095 required an optimal excitation of 610 nm in the Aβ plaques and 660 nm for the tau deposits. This shift in excitation could be attributed to exciton coupling^47^ created by the intermolecular stacking of individual dye molecules along the fibril axis which has been observed in ligand-complexed cryo-EM structures of the amyloids and in multispectral confocal microscopy^41, 48^. Thus, while more work is needed to understand the molecular features of L095 binding, these results suggest that L095 can be used to spectrally differentiate between two related amyloid structures *ex vivo*. More broadly, these findings suggest that *in vitro,* high throughput screens can produce reagents that are useful in *ex vivo* studies to stain interesting features of pathology in human brain samples.

### Analogs of a coumarin chemotype reveal structure-activity relationships

Finally, if the fluoroprobes were binding to discrete site(s) on amyloid fibrils, we expected that a medicinal chemistry campaign might reveal structure-activity relationships (SAR). The alternative hypothesis is that these dyes are simply “sticky”, such that close analogs might have similar or even identical properties. To test these ideas, we focused on the coumarins. Our screening results had shown that these dyes are in the category of pan-fibril binders, meaning that they interact with both WT and P301S fibrils (see above). Thus, we considered that the coumarins might potentially be “sticky”. To assemble an analog set, we searched the MWC Library^21^ to select 24 additional coumarins **(Fig. S5)**. Then, these compounds were tested for interactions with the 26 WT and P301S tau fibril samples in the paDSF-based primary screening platform (**Fig. 6b**). Using three replicates, we found that only four dyes (compounds 15, 16, 22 and 23) bound to most all of the WT and P301S fibrils (**Extended Data 5**). This steep SAR (only 4/24 actives) supports the idea that the dye-binding site(s) have discrete features, rather than being non-specific or “sticky”. Moreover, it tentatively appears that the tertiary amine pendant from the core coumarin might be important for binding (**Fig 6c**), as observed in compounds 15 and 22. Finally, we noted that a subset of the analogs had more restrictive activity, only labelling either WT or P301S tau fibrils in the presence of specific inducers (**Fig 6d**), suggesting that selectively might be obtained through future synthetic efforts. While preliminary, these findings suggest that medicinal chemistry efforts can be used to further optimize validated chemical series from the screening workflow.

**Figure 6.**
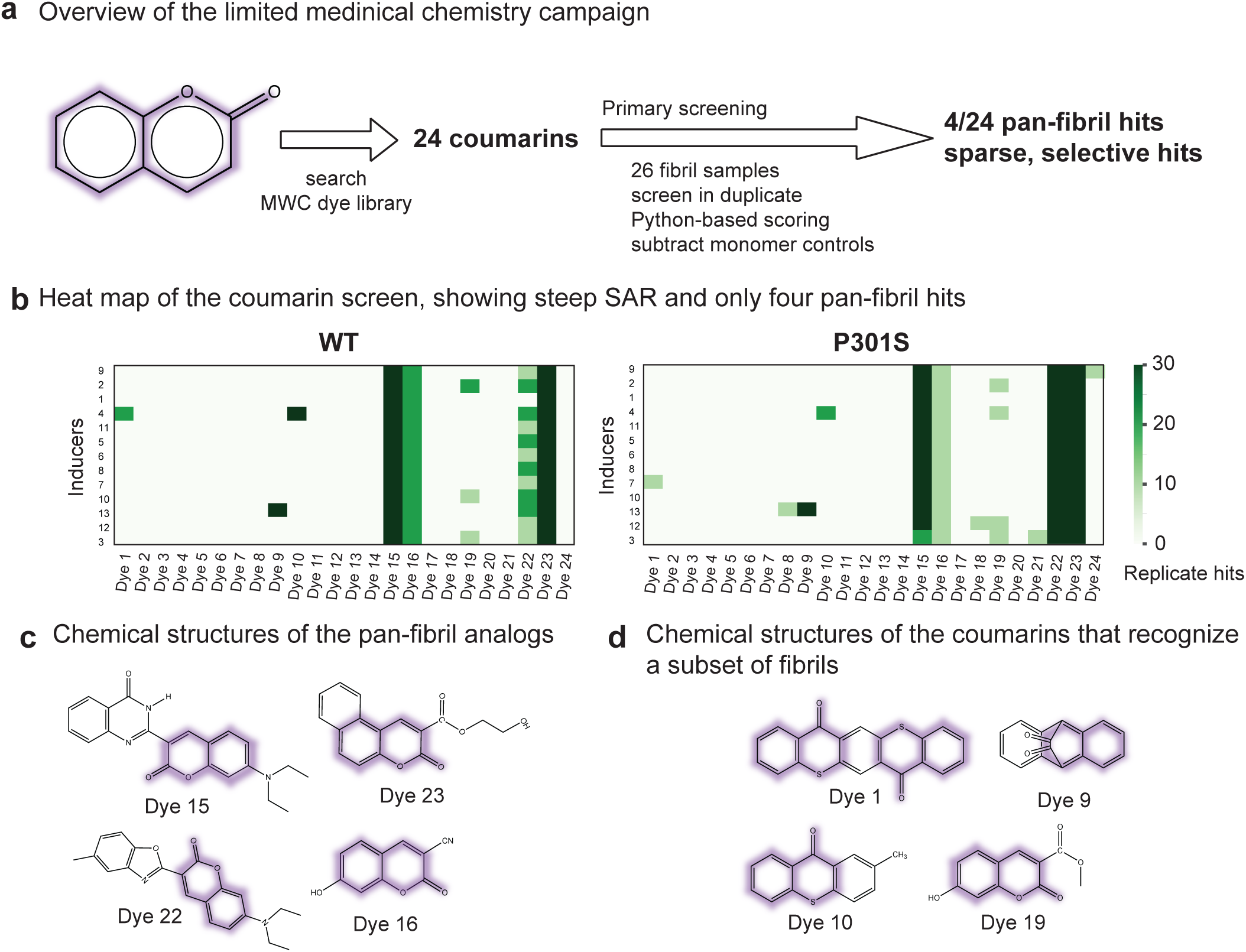
Analogs of the coumarin scaffold include both pan-fibril binding and potentially selective fluoroprobes. (a) Overview of the limited medicinal chemistry campaign. Twenty-four coumarins from the Max A. Weaver Collection (MWC) were screened against 26 fibril samples by paDSF using a pipeline that parallels Figure 1, except that three biological replicates were used. After triage and removal of dyes that bound tau monomer, only 4 coumarins were identified that bind both WT and P301S fibrils. (b) Heat maps of the screening results, showing that most of the analogs failed to recognize either the WT or P301S tau fibril samples (white); whereas a subset produced reproducible signal (green). Here, the top Python score was 30 (10 for each replicate). (b) The chemical structures of the coumarins that bind to both WT and P301S fibrils, suggesting that they are pan-binders. (c) Chemical structures of coumarins with activity against at least one WT or P301S tau fibril conformer. The coumarin scaffold is highlighted in purple.

## Discussion

Fluorescent reporters of tau fibril conformation could be used to dissect the molecular underpinnings of tauopathy and to create diagnostics, such as starting points for positron emission tomography (PET) imaging probes^49, 50^. However, most current fluoroprobes, such as ThT, bind many amyloids and do not discriminate between polymorphs. While other groups are creating analogs of known chemotypes to improve selectivity, such as fluoroprobes that discriminate between oligomers and fibrils^45, 51–54^, we sought to supplement those efforts with a large-scale, unbiased screen of diverse organic dyes. Accordingly, we established a paDSF-based screening platform, coupled with the Aurora collection of dyes, to uncover tau fibril-binding fluoroprobes (see **Figs. 1** and **2**). This screen had a surprisingly low hit rate (10/306 total; ∼3%), suggesting that dye binding involves discrete binding sites instead of non-specific contacts. Indeed, a limited medicinal chemistry campaign, using 24 analogs of the coumarin chemotype, generally supported this idea (see **Fig 6**). This model is also re-enforced by classic work to define the binding sites of ThT and Congo Red^11, 13^, and recent cryo-EM studies, which have shown that the PET probe GTP-1 and the amyloid ligand, epigallocatechin (ECGC), bind to specific sites on tau fibrils^48, 55^. Importantly, fibril polymorphs are known to differ dramatically in their folds and availability of potential pockets^56^, supporting the idea that organic dyes seem well suited to exploit the structural differences between tau fibril cores.

The major goal of this study was to demonstrate whether a paDSF based workflow (see **Fig 1**), combined with an imaging-based EMBER platform (see **Fig 3**) and “ThT-like” kinetic aggregation assays (see **Fig 4**), could yield fluoroprobes that are selective for subsets of tau polymorphs. Because we used synthetic tau fibrils that we do not assume to resemble patient-derived folds, we were less concerned with the exact identity of the “hits” than with the performance of the screening workflow and discriminatory potential of the triage steps. Using 26 fibril samples and 306 Aurora dyes, we confirmed 27/306 (∼9%) initial fluoroprobes that bind at least one of the fibril samples by paDSF. Then, these dyes were further triaged using EMBER to focus on the 10/306 (∼3%) most promising and 7/10 (∼2% overall) were also found to be active in kinetic aggregation assays. Overall, the assay performance was good, with a final, validated hit rate that might be expected for screening campaign carried out with a focused chemical library. However, we also noted that there was relatively poor reproducibility between biological replicates (< 50%), with many dyes scoring well in one experiment and poorly in the other (see **Fig 1**). Thus, one lesson learned from these experiments is that it might be important to leverage biological replicates early in the primary/secondary screening. Another important lesson was that multiple, independent secondary assays were needed to focus on the most robust fluoroprobes for specific uses. Here, the EMBER assay seemed particularly good at identifying dyes that discriminate between polymorphs, while the ThT-like assays provided initial estimates of whether the fibril structure forms early or late in the aggregation process. At this stage, we favor using the two orthogonal secondary assays independently to identify reagents for specific uses. For example, MWC034 might have been de-prioritized by EMBER (see **Fig 3**), but it was a robust reporter in the “ThT-like” kinetic assays (see **Fig 4**). Our hope is that this workflow could inspire screening of patient-derived fibrils, which will likely require further miniaturization. Alternatively, many groups are pursuing ways of generating disease-relevant synthetic fibrils *in vitro*^34, 57, 58^ that recapitulate disease polymorphs.

Another tentative goal of this work was to show whether high throughput approaches might be useful in revealing new or under-explored chemotypes in diverse libraries. Indeed, the validated “hits” included coumarin and polymethine scaffolds. Coumarin-containing probes are documented to have amyloid-binding properties^59–61^, but, to the best of our knowledge, none have been shown to bind tau fibrils. Likewise, polymethine and cyanine dyes, particularly those with relatively short polymethine chains^62^, have only been sporadically identified as having amyloid binding properties^63^, but remain underexplored. Perhaps most importantly, two dyes from the Max A. Weaver library were identified that have not yet (to our knowledge) been reported as binding to amyloids. Thus, we conclude that high throughput approaches might be a good complement to the field’s ongoing efforts to develop polymorph-selective fluoroprobes, with the possibility of identifying new chemotypes.

## Methods

### Ethical Statement

All animals were housed in a facility accredited by the Association for Assessment and Accreditation of Laboratory Animal Care International, in accordance with the Guide for the Care and Use of Laboratory Animals. All procedures for animal use were approved by the University of California San Francisco’s (UCSF’s) Institutional Animal Care and Use Committee. The de-identified, human post-mortem tissues were determined to be exempt from institutional review board approval in accordance with UCSF policy and sourced from UCSF’s Neurodegenerative Disease Brain Bank (https://memory.ucsf.edu/neurodegenerative-disease-brain-bank).

### Purification of tau proteins

*Escherichia coli* BL21 Rosetta 2 (DE3) cells were transformed with pEC135 (WT 0N4R tau) or pEC14<u>6</u> (Tau P301S 0N4R). Cells were grown to between 0.4 and 0.8 optical density (OD_600_) in Terrific broth (TB) and then induced with 1 mM IPTG for 3 hours at 37 °C. Bacteria were pelleted by centrifugation (4,000 xg) and then resuspended in 1x distilled phosphate buffered saline (DPBS) buffer (Corning, pH 7.4) with 10 mM EDTA, 2 mM MgCl_2_, 1 mM dithiothreitol (DTT), and 1x protease inhibitor tablets (Pierce). Resuspended cells were lysed by sonication and the lysate was clarified by centrifugation at 25, 000 rcf at 4 °C for 30 min. Tau variants were first purified via their N-terminal 6x-His tags by incubating the clarified lysate with cOmplete™ His-Tag Purification Resin (Roche, EDTA-and DTT-compatible) for 1 hour at 4 °C. Bound resin was washed with 1x DPBS buffer (Corning, pH 7.4) with 10 mM EDTA, 2 mM MgCl_2_,1 mM DTT, and 1x protease inhibitor tablets (Pierce) and eluted with 1x DPBS buffer (Corning, pH 7.4) with 300 mM imidazole (pH 7.4), 10 mM EDTA, 2 mM MgCl_2_,1 mM DTT, and 1x protease inhibitor tablets (Pierce). Eluate was dialyzed overnight at 4 °C using “tau buffer”: 1x DPBS buffer (Corning, pH 7.4) with 2 mM MgCl_2_,1 mM DTT. Concentration of purified protein was measured using the BCA assay (Pierce). Then, a reverse phase chromatography step was performed using a Kromasil semipreparative column (250 mm length, 10 mm I.D., C4, 5 µm particle size, 300 Å pore size) equilibrated with 5% acetonitrile. Tau protein was eluted from the column using a 5-50% acetonitrile gradient over the course of 45 minutes. Fractions containing pure, full-length tau were then lyophilized to remove solvent and resuspended in tau buffer and stored at -80 °C.

### Creation and purification of tau fibrils

Aggregation reactions were performed in 1.5 mL Eppendorf tubes for 48 hours at 37 °C with constant agitation at 1200 rpm. All reactions contained 10 µM tau, plus a polyanion inducer (see below) in tau buffer (1x DPBS buffer (Corning, pH 7.4) with 2 mM MgCl_2_,1 mM DTT) at a final volume of 300 µL. Polyanion inducer concentrations were calculated from the midpoint concentration for successful amyloid induction in previous studies^17^ and are found in **Table S1**. All inducer stocks and buffer were freshly prepared each day. After 48 hours, reactions were ultracentrifuged at 103,000 rcf using a tabletop Beckman Optima Max-XP Ultracentrifuge with TLA-55 rotor. Pellets containing tau fibrils were resuspended in tau buffer, tested for positive ThT fluorescence to validate the presence of amyloid, and quantified using A_205_ absorbance on a Nanodrop Microvolume Spectrophotometer (Thermo Fisher).

### paDSF screens

Tau fibrils (at least 100 µg/mL) were screened for binding to 306 members of the Aurora dye library.^20^ Briefly, screens were performed in 384-well plate format (Axygen) using a qTower real-time thermal cycler (Analytik Jena). Fibril stocks (2 µL) in tau buffer were added to individual wells, followed by addition of dye (8 µL; final concentration between 0.5 to 50 µM, depending on the optimal value determined previously^20^). These mixtures were incubated at room temperature for 10 minutes. Three control screens were also performed. The first samples contained dye, but no fibril (termed the “no protein control”). The second contained dye and polyanion (termed the “polyanion only control”). The third samples contained monomeric tau (1 µM) without the polyanion inducer (termed the “tau monomer control”). Fluorescence was monitored for six excitation/emission wavelength pairs: 470 nm/520 nm (FAM), 515 nm/545 nm (JOE), 535 nm/580 nm (TAMRA), 565 nm/605 nm (ROX), 630 nm/670 nm (Cy5), and 660 nm/705 nm (Cy 5.5). Fluorescence was measured as a function of temperature, with a temperature ramp of 1 °C increase per cycle from 25 to 95 °C over the course of one hour.

### paDSF data analysis

Potential candidates for dye-fibril interaction were identified based on fluorescence profiles. Specifically, we sought to identify samples with high initial fluorescence, followed by diminished signal intensity as temperature increased (presumably due to thermal melting or re-arrangement of the binding site(s)). To identify the top performing dyes, a Python-based scoring function was used, and this scoring function is available in full ***(Extended Data 1)***. Briefly, the scoring function assigns a score from 0 to 10 and scores were assigned as follows:

| Score | Max. fluorescence (RFU) | Min. fluorescence (RFU) | Δfluorescence (fibril-buffer) (RFU) | Temp. Max. fluorescence ($^{\circ}\text{C}$) |
| --- | --- | --- | --- | --- |
| 10 | >1000 | >-1000 | >2500 | <55 |
| 8 | >1000 | >-2000 | 1500<x<2500 | <55 |
| 5 | >1000 | >-2000 | 1000<x<1500 | <55 |
| 3 | >2000 | >-2000 | >2000 | 55<x<70 |
| 1 | >2000 | >-2000 | >2000 | >70 |

All other fluorescence profiles received a score of 0.

All dye-fibril combinations that scored a 10 were manually inspected to remove false positives. The most common artifact resulted from bright dyes, in which small pipetting/concentration differences seemed to result in large, but misleading, apparent RFU differences. Second-generation coumarin analogs (see **Fig 5**) were screened in the same manner with a concentration of 0.5 µM dye and scored manually as either ‘hit’ or “non-hit”. For each of three replicates, hits were assigned a value of 1 (*i.e.,* a score of 3 indicates the second-generation dye was a hit in three independent screening experiments).

### EMBER confocal microscopy

The 27 dyes hits that passed initial screening steps described above were validated for direct binding to fibrils via fluorescence confocal microscopy imaging. For initial imaging, WT and P301S fibrils induced with heparin, sodium alginate, and sodium hexametaphosphate (six fibril types) were deposited in individual wells of a Corning™ BioCoat™ 384-Well, Collagen Type I-Treated, Flat-Bottom Microplate. The plate was centrifuged at 50 rcf to deposit fibrils on the bottom of the plate. Dyes were diluted to screening concentrations (0.5 to 50 µM) in tau buffer (1x DPBS buffer (Corning, pH 7.4) with 2 mM MgCl_2_ and 1 mM DTT), centrifuged for 10 minutes at 17,000 rcf to remove dye aggregates, and added to fibril-containing wells. If dye binding to fibril occurred, fibrils were imaged using our spectral confocal microscopy method, called EMBER imaging^22^. Fibrils stained with each active dye were imaged using a Leica Microsystems SP8 confocal microscope equipped with a 40x water-immersion lens (1.1 NA) and utilized a white light and 405 nm lasers along with a HyD detector to capture images at a resolution of 512x512 pixels at 1x zoom. The optical plane was autofocused with the highest sensitivity setting for each field of view. To minimize background noise, the LightGate was set between 0.5-18 ns. Initially, 110 images were captured using the Λ/λ-scan mode with wavelengths ranging from 470 nm to 670 nm at 20 nm intervals. The emission detection started at 10 nm above the given excitation wavelength and concluded at 780 nm within a 20-nm window. For instance, for the 470-nm excitation, images were collected from 480 nm to 780 nm at 20-nm intervals. Subsequently, in the λ-scan mode, 18 more images were captured at 405-nm excitation with emission detection intervals of 20 nm, ranging from 420 nm to 780 nm. These results are available in **Extended Data S6**.

### EMBER conformational analysis of tau fibrils

The EMBER analytical pipeline to discriminate conformational strains of fibrils is described previously^22^. In brief, we developed a set of MATLAB scripts to process raw fluorescence images and to segment fibril particles from the background. The scripts yielded a particle-resolution EMBER profile for all recognized particles in each confocal experiment. Subsequently, we normalized each identified particle-resolution EMBER spectra and concatenated EMBER profiles from both WT and P301S 0N4R tau fibrils corresponding to tested polyanions and specific dye. The data was then subjected to dimension reduction using Principal Component Analysis (*pca*). To quantify the discrimination power of a specific dye against WT and P301S 0N4R tau fibrils, we implemented a quadratic fit discriminant analysis classifier (*fitcdiscr*). The accuracy scores from this analysis served as the discrimination score, which were averaged across polyanions per specific dye to calculate an overall discrimination score (see **Fig 3**).

### Calculation of molecular similarities

Tanimoto similarities of all pairwise dye hit combinations were calculated using the RDkit Python package (https://www.rdkit.org/docs/index.html and https://github.com/rdkit/rdkit).

### Real-time kinetic fibril detection assays (“ThT-like” assay)

Aggregation assays were performed in 384-well microplates (Corning 4511) coated with 0.01% Triton-X detergent. WT or P301S tau (10 µM) was combined with 50 µM dye (L016, L017, L033, L063, MWC034, MWC061, L079, L089, and L095), 5 µM dye (L031), or 10 µM Thioflavin T (ThT) and freshly prepared polyanion inducer (see **Table S1** for concentrations) in individual wells to a final volume of 20 µL in tau buffer (1x DPBS buffer (Corning, pH 7.4) with 2 mM MgCl_2_,1 mM DTT). Aggregation reactions were carried out in a SpectraMax M5 microplate reader (Molecular Devices) at 37 °C with continuous shaking for 48 hours. Fluorescence was monitored for four excitation/emission wavelength pairs: 444 nm/482 nm, cutoff 475 nm; 470nm/520 nm, cutoff 515 nm; 515 nm/545 nm, cutoff 530 nm; 565 nm/605 nm, cutoff 590 nm. Each reaction was performed in triplicate. To facilitate comparison of dyes between different reactions, RFU values were normalized between 0 and 1. We did not test all of the tau fibril samples in these experiments and, rather, focused only on the samples that had shown the most promising results from the EMBER studies.

### Ex vivo imaging of amyloid plaques in mouse brain

Samples were obtained from APP23 or 5xFAD mice (see **Fig. S4**). To reduce the autofluorescence in the brain tissue, we photobleached the sections for up to 48 h in a cold room using a multispectral LED array.^64^ The sections were then deparaffinized, and subjected to hydrolytic autoclaving at 105 °C for 20 min in citrate buffer (Sigma, C9999). Following blocking with 10% normal goat serum (Vector laboratories, S-1000), sections were incubated with primary antibody anti-β-amyloid, 17-24 Antibody clone 4G8 (Biolegend 800702) 1:1000, overnight at room temp. After washing, sections were incubated in secondary antibody Alexa Fluor goat anti-mouse 405 (Thermo Fisher A48255) 1:500 for 120 minutes at room temp. Sections were then washed in PBS and incubated in L095 (50 µM) for 20 minutes and then rinsed with DI water and cover slipped using Permafluor aqueous mounting medium (Thermo Scientific, TA030FM). Brain sections were imaged using a Leica Microsystems SP8 confocal microscope equipped with a 40x water-immersion lens (1.1 NA) and utilized a white light and 405 nm lasers along with a HyD detector to capture images at a resolution of 512x512 pixels at 2x zoom. To minimize background noise, the LightGate was set between 0.5–18 ns.

### Imaging of human AD patient samples

Samples for immunofluorescence were formalin fixed paraffin embedded and cut at 8 µm. Slides were deparaffinized and subjected to hydrolytic autoclaving at 121 °C for 10 min in citrate buffer (Sigma, C9999). Following blocking with 10% normal goat serum (Vector laboratories, S-1000), sections were incubated with primary antibody AT8 (1:250; Thermo Fisher MN1020) or anti-tau phospho-S396 antibody [EPR2731] (abcam ab109390) 1:500 and 4G8 (1:1000; Biolegend 800702) overnight at room temp. After washing, sections were incubated in secondary antibodies Alexa Fluor goat anti-rabbit 488 (Thermo Fisher A11008) and Alexa Fluor goat anti-mouse 405 (Thermo Fisher A21235) both 1:500 for 120 minutes at room temp. L095 was used as above. Sections were then washed with DI water and coverslipped using Permafluor aqueous mounting medium (Thermo Scientific TA030FM). Brain sections were imaged using a Leica Microsystems SP8 confocal microscope equipped with a 40x water-immersion lens (1.1 NA) and utilized a white light and 405 nm lasers along with a HyD detector to capture images at a resolution of 512x512 pixels at 2x zoom. To minimize background noise, the LightGate was set between 0.5–18 ns.

## Data Availability

The Aurora library is found in **Extended Data 1** (https://doi.org/10.5281/zenodo.13357986). The uncropped blots are available in **Extended Data 2** (https://doi.org/10.5281/zenodo.13357986). The full primary paDSF screening results and Python code are available as **Extended Data 3** (https://doi.org/10.5281/zenodo.13357986). The kinetic assay data is available in **Extended Data 4** (https://doi.org/10.5281/zenodo.13357986). The screening data for the coumarin analogs is found in **Extended Data 5** (https://doi.org/10.5281/zenodo.13357986). The raw EMBER data is available in **Extended Data 6**(https://datadryad.org/stash/share/YS5EBV9EomjU_vCXUm7-1Wzq23WTkVTyVUg3wuRgBA4 or doi:10.5061/dryad.s4mw6m9g4). All other data is available in the Supplementary Information.

## Acknowledgements

We thank Taia Wu (UCSF) for technical advice on paDSF and Eric Greene (UCSF) for assistance with Python. We thank Wyatt Powell for comments on the manuscript. This work was supported by grants from Tau Consortium (to J.E.G.) and the National Institutes of Health: F32AG076281 to E.C.C.), R01GM141299 (J.E.G. and N.R.V.), P01AG002132 (C.C. and W.F.D.), RF1NS133651 (C.C.), and GM122603 (W.F.D.). H.Y. was supported by the NIH (K99AG084926). The microtiter plate in Fig. 1a was adapted from a biorender.com icon. Tissue samples were supplied by the London Neurodegenerative Diseases Brain Bank (King’s College London, England), which receives funding from the Medical Research Council UK and through the Brains for Dementia Research Project (jointly funded by the Alzheimer’s Society and Alzheimer’s Research UK).

## Author Contributions

E.C. and J.E.G. conceptualized the studies. E.C., H.Y., J.G.J., A.O. and A.F.C. performed experiments and interpreted results. A.Y., K.M.M., Z.M. and N. R.V. provided reagents. N.R.V., W.F.D., D.A.M., C.C. and J.E.G. provided supervision and funding. E.C. and J.E.G. drafted and edited the manuscript. All authors edited the manuscript.

