## Supplemental Information for "Methods for high throughput discovery of fluoroprobes that recognize tau fibril polymorphs"

| Number | Polyanion name | CAS Number | Supplier | Effective Induction Concentration (μg/mL) |
|---|---|---|---|---|
| 1 | Heparin Sodium | 9041-08-1 | Santa Cruz Biotechnology | 100 |
| 2 | Sodium Polyphosphate | 10361-03-2 | | 1000 |
| 3 | Poly-L-Glutamate sodium salt | 26247-79-0 | Sigma | 500 |
| 4 | Fondaparinux sodium | 114870-03-0 | Sigma | 500 |
| 5 | Nadroparin calcium | 37270-89-6 | | 250 |
| 6 | Sodium alginate | 9005-38-3 | Sigma | 500 |
| 7 | Sodium hexametaphosphate | 68915-31-1 | | 500 |
| 8 | Polystyrene sulfonate | 25704-18-1 | | 62.5 |
| 9 | Poly(A) | 26763-19-9 | Sigma | 500 |
| 10 | Sodium Tripolyphosphate | 7758-29-4 | | 2000 |
| 11 | Chondroitin sulfate A sodium salt | 39455-18-0 | Sigma | 250 |
| 12 | Diadenosine pentaphosphate | 75522-97-3 | Sigma | 500 |
| 13 | Dermatan sulfate and oversulfated chondroitin sulfate | NA | Sigma | 125 |

**Table S1. Summary of polyanion inducers used to generate WT and P301S tau fibril polymorphs used in this study.**

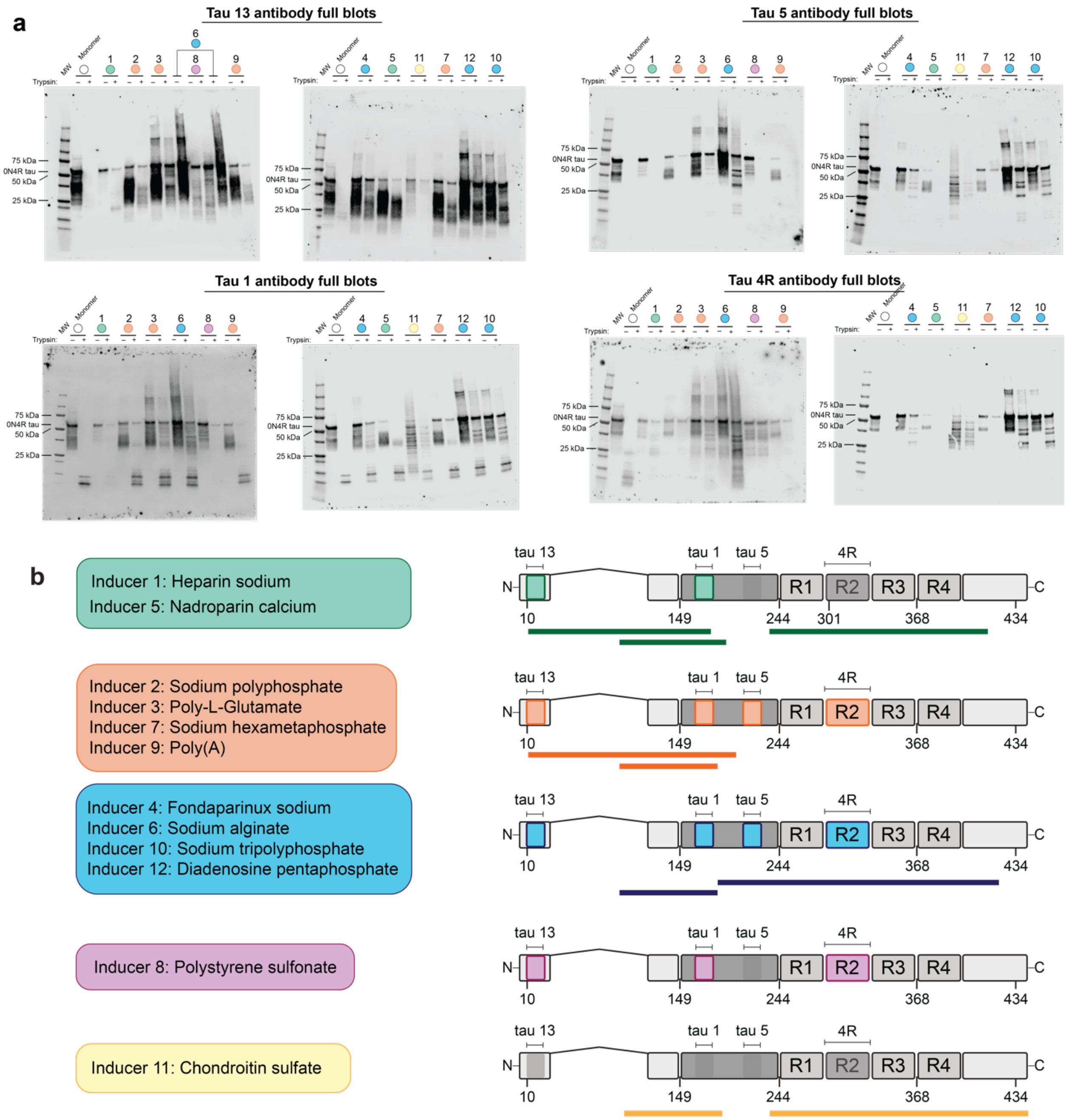

**Figure S1. Inducer-generated P301S tau fibrils also adopt diverse conformations that may be distinct from WT tau fibrils. (a)** Western blots with antibodies recognizing four different tau epitopes from limited proteolysis experiments performed with Promega sequencing grade trypsin at a fibril (or monomer):trypsin ratio 500:1 for 60 min. at 37 °C, as performed previously for WT tau in ref. 29 . The fibrils are generally resistant to trypsin cleavage compared to monomer, but yield distinct protease-resistant fragments that suggest considerable fibril conformational diversity among different inducers. **(b)** (Left) Categorization of polyanion-inducer generated fibrils that possess similar—but not necessarily identical—proteolysis patterns. (Right) Mapping of protease-resistant fragments to the 0N4R Tau P301S primary structure based on resistance of tau epitopes to cleavage for each category. Protease-resistant epitopes are shown in color with lines below the schematic representing general putative protease-resistant fragments observed within each category (not necessarily exact fragments for each inducer-generated fibril).

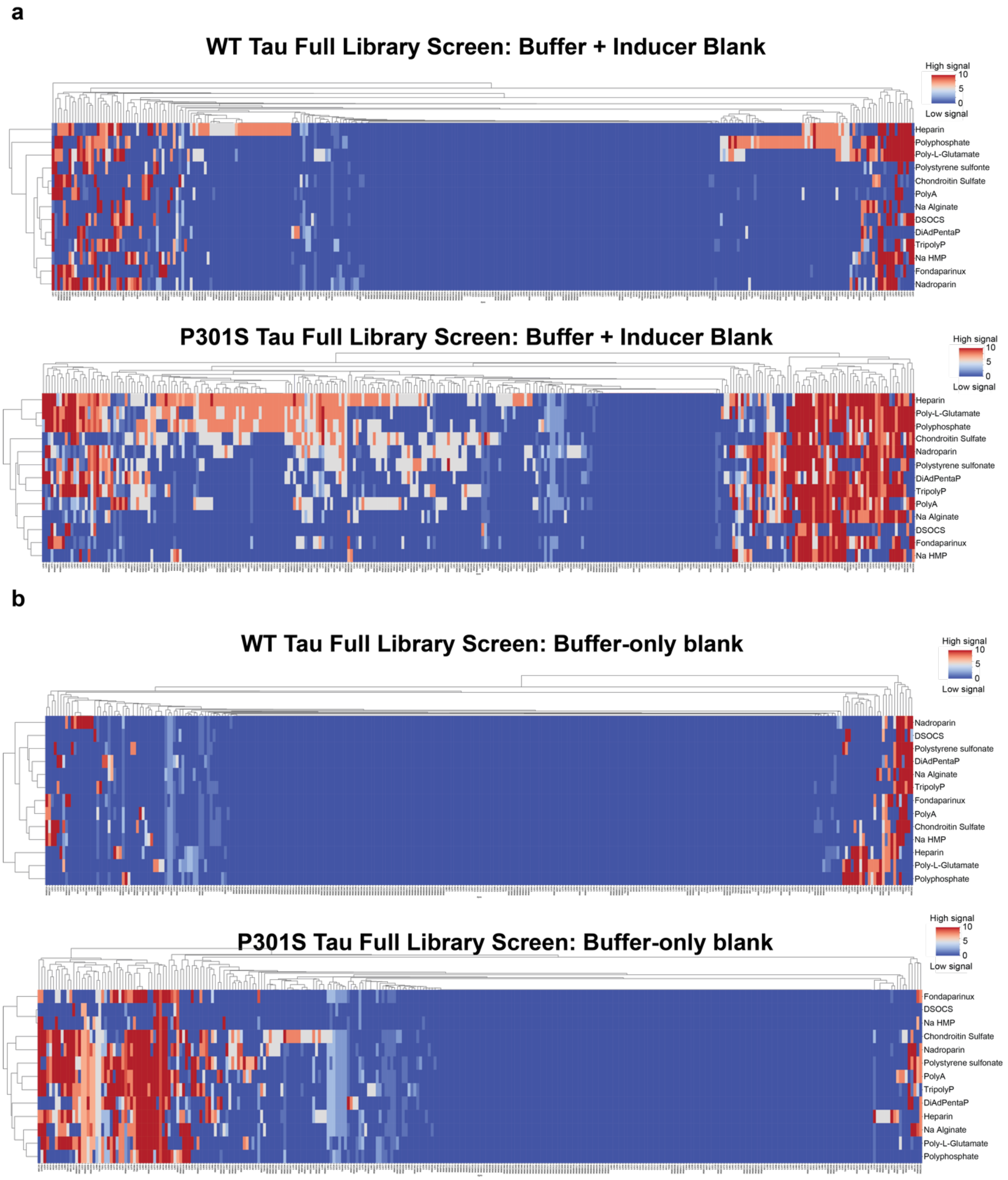

**Figure S2. Heat maps displaying full Aurora library screening data analyzed with different buffer blanks. (a)** Heat maps depicting scores generated from Python scoring function (see Extended data 2) of all Aurora library fluoroprobes incubated with 13 polyanion-induced WT tau fibrils (top) and P301S tau fibrils (bottom) using tau buffer (see methods) supplemented with the screening concentration (Table S1) of each polyanion **(b)** Heat maps depicting scores generated from Python scoring function (see Extended data 2) of all Aurora library fluoroprobes incubated with 13 polyanion-induced WT tau fibrils (top) and P301S tau fibrils (bottom) using tau buffer (see methods) alone.

| Dyes Tested for fibril binding using Confocal Microscopy | Binding Observed? |
|---|---|
| C018 | no |
| L009 | no* |
| L016 | yes |
| L017 | yes |
| L018 | no |
| L031 | yes |
| L033 | yes |
| L062 | no |
| L063 | yes |
| L073 | no |
| L077 | no |
| L079 | yes |
| L080 | no |
| L081 | no |
| L089 | yes |
| L095 | yes |
| L105 | no |
| MWA010 | no |
| MWA013 | no |
| MWC002 | no |
| MWC024 | no* |
| MWC027 | no |
| MWC034 | yes |
| MWC061 | yes |
| MWE07 | no |
| MWE08 | no |
| MWF03 | no |

*some potential binding observed but poor dye photophysics

**Table S2. Summary of 27 Aurora library fluoroprobes tested for direct binding to tau fibrils using confocal microscopy.** Yellow highlight indicates that a fluoroprobe is one of the ten final validated hits.

Dye hit L031 and Thioflavin T curves track closely across multiple inducers, perhaps due to a shared molecular rotor binding mode

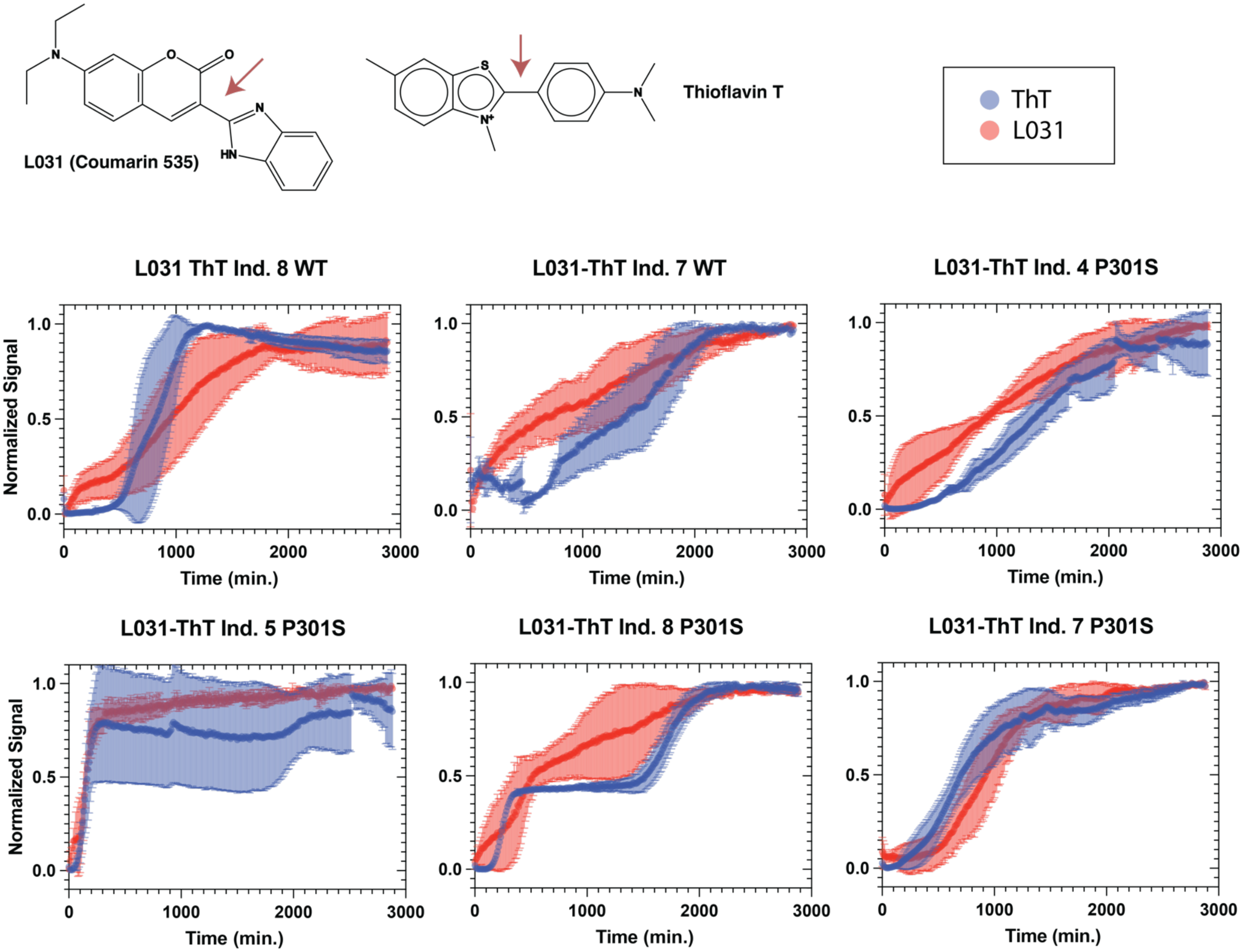

**Figure S3. Fluoroprobe L031 and Thioflavin T (ThT) curves track closely across multiple inducers, perhaps due to a shared molecular rotor binding mode.** Kinetic aggregation experiments performed with both WT and P301S tau and diverse polyanions reveal that the fluorescence profiles of dye L031 resemble those of ThT across multiple inducers. Thus, L031, which shares a molecular rotor architecture with ThT, may possess similar molecular recognition properties to (i.e., bind to similar binding sites during fibril formation) based on this close correspondence between their aggregation profiles. Both signals were monitored in the ThT channel (ex: 444, em: 482), n=3.

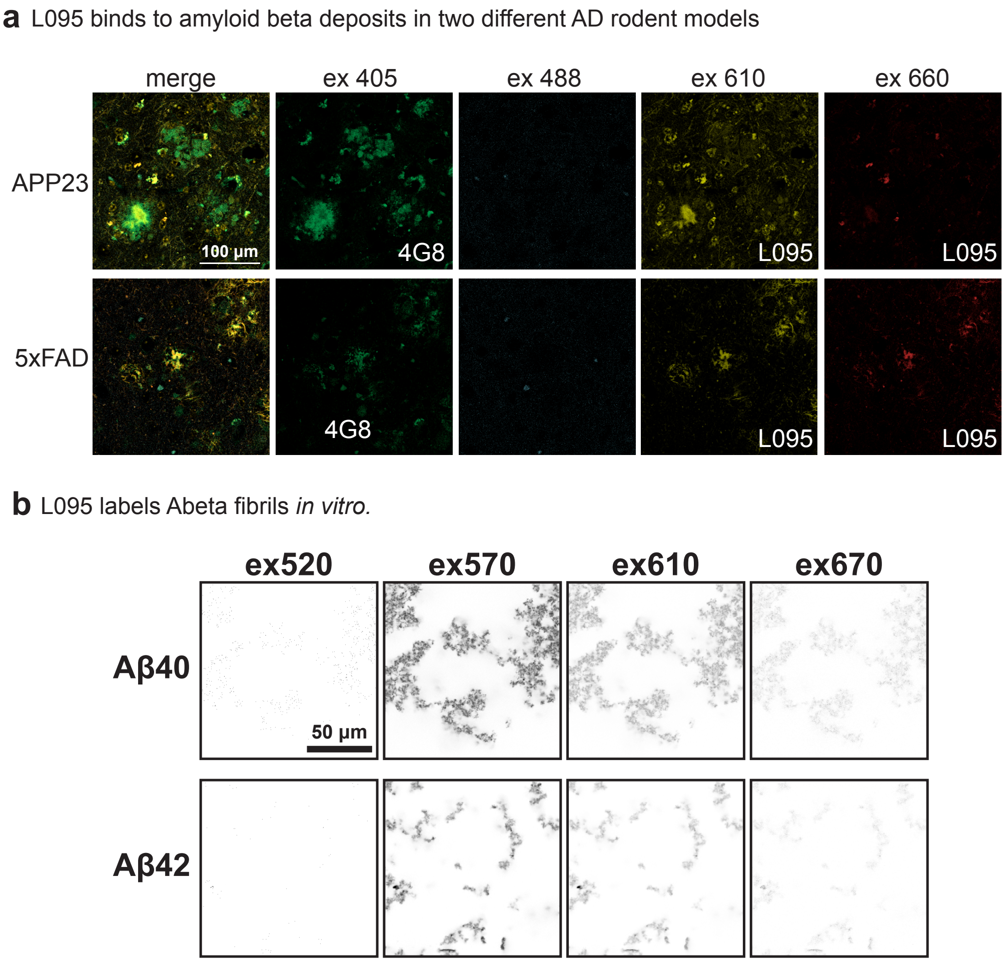

**Figure S4. Control experiments show that dye L095 also binds to amyloid-β.** (a) Staining of histology slides from two transgenic mouse models of Alzheimer's disease, showing overlap of dye L095 and an anti-amyloid-β antibody (4G8). Scale bar = 100 µm. (b) Binding of L095 to recombinant Aβ 1–40 and Aβ 1–42 fibrils, as measured by EMBER.

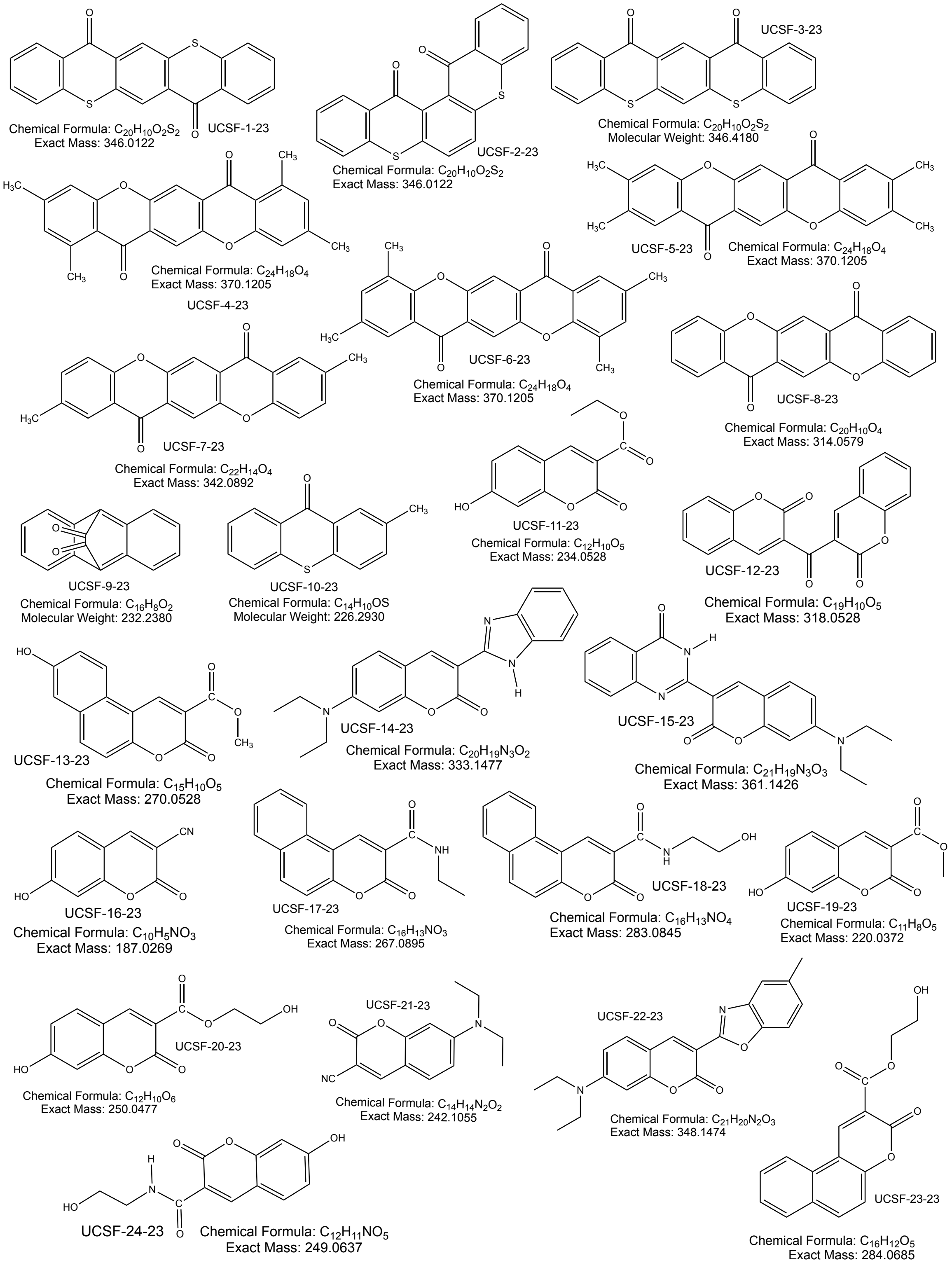

**Figure S5. Full chemical structures and formulas with molecular weights (in g/mol) for all coumarin-analog fluoroprobes discovered in a structural homology search of the Max Weaver Dye Library.** All compounds were screened with the 26 polyanion-induced fibrils in the second-generation paDSF screens.

| Cohort | Paitent ID # | Sex | Age | NPDx | Mutation | APOE genotype | CERAD score | Braak stage | Brain region | Source |
|---|---|---|---|---|---|---|---|---|---|---|
| AD 1 | A2201 P2926B4 | M | 82 | AD | – | – | C3 | VI | Occipital | UCSF IND |
| AD 2 | BBN13829 A029/98x30 | M | 42 | AD | Δ4 PSEN1 | 3/3 | C3 | VI | Temporal cortex | King's College London (UK) |
| AD 3 | BBN13932 A0258/94 | F | 55 | AD | V717I APP (London) | 3/4 | C3 | VI | Temporal cortex | King's College London (UK) |

**Table S3. Sources of postmortem human brain tissue samples.**
